# Disparity in the Distribution of Suitable Habitat: A Metric for Capturing and Tracking Responses to Urbanized Landscapes

**DOI:** 10.1101/2025.10.15.682594

**Authors:** Joscha Beninde

## Abstract

Global species observations from community science platforms offer an unprecedented opportunity to analyze large-scale biodiversity patterns. This holds great promise for developing data-driven metrics to measure species’ responses to landscape-level environmental changes, essential for ameliorating the ongoing biodiversity crisis. A primary cause of declines in local, native species is the expansion and intensification of urbanized landscapes. A species’ response to urbanization can remain consistent across spatial and temporal scales. However, city-specific factors, such as land-use dynamics or local biotic interactions, can profoundly influence species and cause shifts in responses. This underscores the importance of measuring responses to urbanized landscapes at the local population level in a way that facilitates spatial and temporal comparisons. I propose utilizing the receiver operating characteristic (ROC) and its area under the curve (AUC), a statistical classifier, in a novel way for this purpose. Specifically, the disparity in suitable habitat between urban and adjacent non-urban areas is summarized by a single metric, HD_URBAN_, which facilitates population-level comparisons across time and space. I apply it to geographic predictions of the realized niche of 1,023 species from a highly urbanized landscape in California, USA. Additionally, modeling virtual species for each of the urban response types introduced by Blair’s seminal 1996 study provides place-based, expected HD_URBAN_ scores for urban avoider, urban utilizer, and urban dweller species. Validation efforts demonstrate accurate predictions of expected urban responses of real species in three scenarios. First, following a route-of-introduction hypothesis, non-native species exhibit significantly higher HD_URBAN_ than native species. Second, assessing the robustness of urban responses in space and time, HD_URBAN_ is significantly correlated to the urban response of the same 38 bird species from Blair’s study, conducted 450 km apart and 25 years earlier. Third, testing for sensitivity to temporal shifts within species, HD_URBAN_ decreases significantly in a species displaced from urban areas due to negative interactions with a recently introduced, congeneric species. In summary, HD_URBAN_ is a versatile metric that provides a data-driven way to compare species responses, supporting future research into the patterns and processes underlying spatial and temporal variation in responses to urbanized landscapes.

## INTRODUCTION

Species occurrence records are increasing rapidly and becoming available through openly accessible community science platforms, such as iNaturalist (Di Cecco et al., 2021) and eBird (Sullivan et al., 2009), providing an unprecedented opportunity to examine large-scale patterns of biodiversity. This holds great promise for developing data-driven metrics quantifying species’ responses to landscape and environmental changes. These metrics can be utilized to understand and mitigate the ongoing biodiversity crisis (Devictor and Bensaude-Vincent 2016, Moll et al. 2019, Bayraktarov et al. 2019). One of the primary causes of local species declines is the expansion and intensification of urbanized and agricultural landscapes (Grimm et al. 2008, Newbold et al. 2015, 2020, Cooper et al. 2021). In urbanized landscapes, patterns in species distribution, abundance, or density are traditionally loosely categorized as a function of urban land use tolerance or association (McDonnell and Pickett 1990, Blair 1996, McKinney 2002, McDonnell and Hahs 2008, 2015, Fischer et al. 2015). In a seminal study published in this journal, Blair (1996) classified bird species responses using a three-tiered approach. He defined *urban avoiders* as being sensitive to urbanization, with peak densities in wildland areas; suburban adaptable species had the highest densities at intermediate levels of urbanization, and urban exploiters had density peaks in urban areas. Considerable further scrutiny (McKinney 2002, Fischer et al. 2015, McDonnell and Hahs 2015) has altered Blair’s typology slightly, and species are now more commonly referred to as urban avoiders, urban utilizers, and urban dwellers (Fischer et al. 2015, Callaghan et al. 2020a). Due to their ease of comprehension and intuitive nature, these response types have largely replaced terms typically used to describe distribution patterns of species in other landscapes, such as resident, short- and long-distance migrant, nomad, or invader (McDonnell and Hahs 2015). Whether this shift in terminology is warranted or not, the three-tiered typology is ubiquitously applied in research and conservation, often without quantitative justification, thereby precluding efforts to develop continuous and comparable metrics of urban responses that can be used more widely.

Data-driven efforts that quantify the magnitude of responses often focus on only a few species, a single taxonomic group, or a single urban landscape (Callaghan et al. 2019; Moll et al. 2020; Fidino et al. 2021; Kuussaari et al. 2021). More comprehensive efforts are emerging (Callaghan et al., 2020a, 2020b; Curti et al., 2023), and quantify responses by summarizing continuous measures of urbanization at species presence locations, e.g., using night-time lights (Callaghan et al. 2020a, 2020b) or PCA-based summaries of urban intensity (Curti et al. 2023). While these approaches provide continuous measures of urban responses and allow for relative species comparisons within the scope of the study, variation in the underlying scale of measurements makes direct comparisons with other species or landscapes challenging or impossible. Moreover, focusing solely on the environmental conditions observed at species presence locations may confound inferences of landscape-level impacts when suitable, but unsampled locations, or the prevalence of environmental conditions within a given landscape, are omitted. Species distribution modeling can directly address these issues by correcting for spatial observation bias and by geographically projecting suitable habitat across the entire study area as a function of environmental conditions (Merow et al. 2013, Valavi et al. 2022; Grether et al. 2024). After accomplishing this, an appropriate metric could contextualize measurements of urban responses on a standardized and readily interpretable scale, enabling comparisons of responses beyond specific taxonomic and spatial contexts.

Metrics of responses to urban areas ideally capture their magnitude accurately while also being sensitive to detect shifts in responses. Overarchingly, urban response types are conceptualized at the species level - as a species’ inherent property. Across species in space and time, urban responses could thus be robust. This has been demonstrated across continental and local scales in avian species, revealing similar urban responses of species across spatial scales (Callaghan et al., 2020a). Documented examples where species’ urban responses change, for example, species’ distributions shifting more towards urban areas due to increasing urban resource utilization (Rutz 2008, Cooper et al. 2021), are less common and frequently restricted to only some populations of a species and in a specific urban landscape. While this may be an exception to the species-level stability of urban responses, it also emphasizes the importance of measuring urban responses locally to detect changes when they occur. For this purpose, I here propose a well-established continuous statistical classifier, the receiver operating characteristic (ROC; Pollack and Decker 1958), and its area under the curve (AUC; Hanley and McNeil 1982; Araújo et al. 2019). This AUC is a continuous summary metric expressed on a scale of 0 to 1 (Hanley and McNeil 1982, Merow et al. 2013). In the species distribution modeling (SDM) literature, AUC is particularly well-known as a metric of model fit, calculated by comparing SDM predictions from presence and absence (or background) locations (Philips 2008, Aruajo et al 2021, Valavi 2021). Adopting this approach, but by comparing SDM predictions from locations in urban and non-urban habitats, AUC is effectively converted into a metric that summarizes the disparity of suitable habitat distribution, or habitat disparity (HD) in short. I here introduce and validate HD_URBAN_ as a metric to quantify the local urban population ecological response of species.

Next to the convenient statistical properties that AUC offers, this approach can be extended easily with a virtual species modeling framework (Hirzel et al. 2001; Meynard et al. 2019) to generate expected HD_URBAN_ values for the three urban response types (urban avoider, urban utilizer, and urban dweller). While computationally demanding, this offers the unique opportunity to directly compare real species to the range of values expected for these urban response types in a focal landscape, and allows for straightforward standardization across landscapes. Here, I calculate HD_URBAN_ for a published SDM dataset of 1,023 species from the Greater Los Angeles metropolitan area, California, USA (Beninde et al. 2023). Due to the steep environmental gradient from urban to wildland habitats in this area (Figure 1a), this dataset is particularly well-suited for evaluating a metric that quantifies a species’ response to urbanized landscapes. The final steps serve to validate HD_URBAN_ as a metric by:

1. Comparing HD_URBAN_ between native and non-native species, expecting higher urban association in non-native than native species, given urban-associated introduction pathways of non-native species (Cadotte et al. 2017, Gaertner et al. 2017).
2. Testing the spatiotemporal robustness of urban responses across species by correlating HD_URBAN_ calculated for contemporary Los Angeles with the urban association ranks observed by Blair in the Bay Area, California, USA (1996), spanning over 25 years and set 450 km apart (Figure 1b).
3. Testing the sensitivity to detect temporal shifts in responses of species, by generating HD_URBAN_ for multiple temporal bins and quantifying the recent decline of the native Western Black Widow, *Latrodectus hesperus*, from urban areas due to the introduction of a congeneric species (Figure 1c; Vetter et al. 2012, Aragon-Traverso 2021).

**Figure 1:**
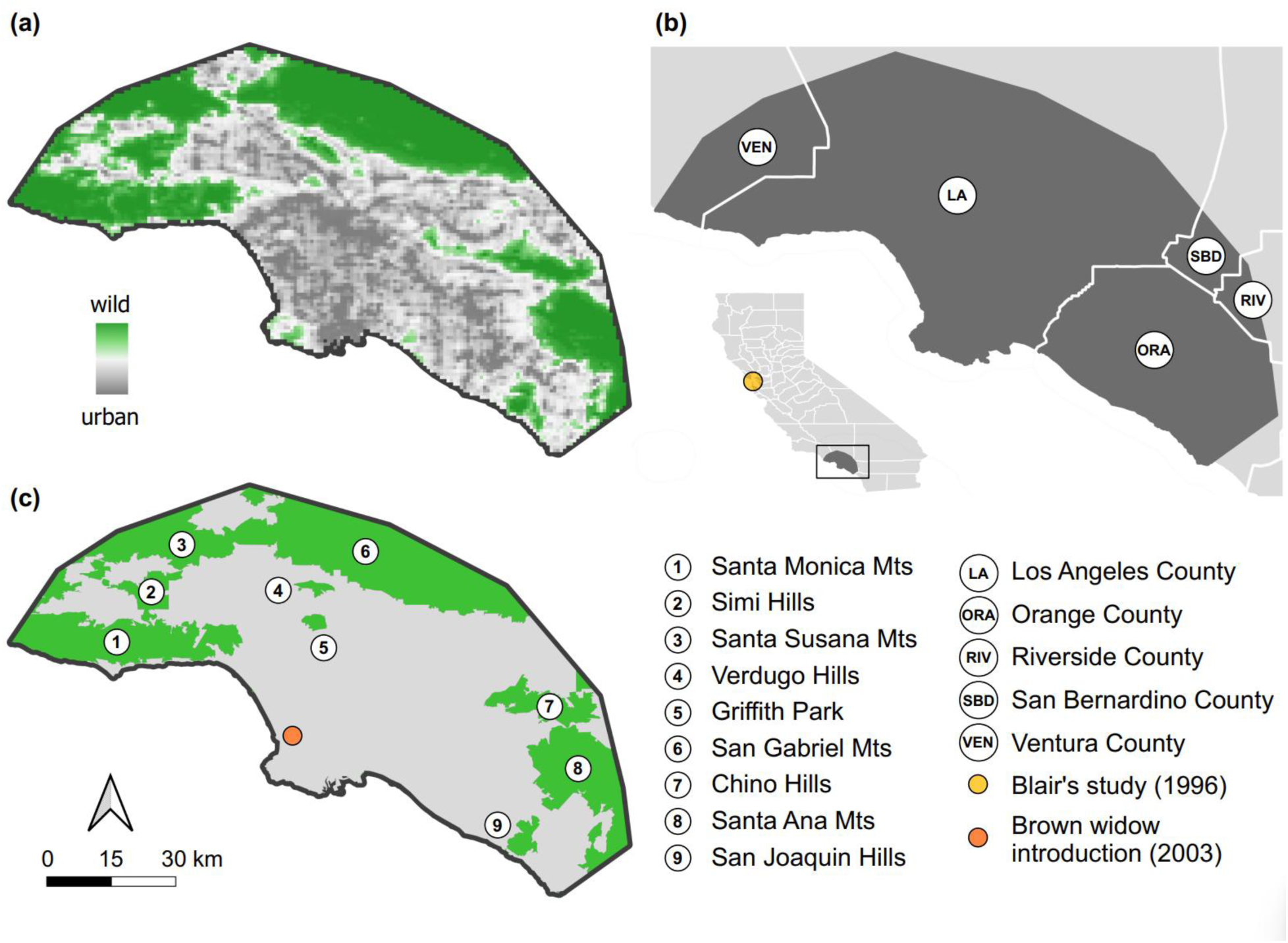
The study area in Greater Los Angeles, illustrating (a) the sharp transition from urban to wildland areas, (b) its location relative to administrative boundaries, the state of California, USA, and Blair’s study location, and (c) the point where the Brown Widow was introduced and the binary land classification into urban and wildland areas used to calculate HD_URBAN_.

## MATERIALS & METHODS

### Overview

This study introduces a metric that measures responses to urbanized landscapes at the local population level, allowing for taxonomic, spatial, and temporal comparisons. It applies the receiver operating characteristic (ROC; Pollack and Decker 1958) in a way that measures the disparity of species’ suitable habitat distribution as a function of a focal (here urban) habitat type by using the area under the ROC curve (AUC; Hanley and McNeil 1982; Araújo et al. 2019). Underlying this is the published habitat suitability dataset for 1,023 species in the Greater Los Angeles region (Beninde et al., 2023). Generated using a species distribution modeling (SDM) framework (also commonly referred to as environmental niche or habitat suitability modeling), the resulting habitat suitability values can be interpreted as a geographic prediction of the realized niche space of species (Merow et al. 2013, Valavi et al. 2022). SDM approaches are informative and frequently utilized in, for example, biodiversity assessments (Riva et al. 2023) or analyses of niche differentiation (Grether et al. 2024), capturing ecologically meaningful results when best-practice modeling standards are met (Araújo et al. 2019). The modeling framework employed by Beninde et al. (2023) meets these standards: it addresses spatial accuracy and biases of observations, temporal resolution, and collinearity of predictor variables, as well as model uncertainty and performance (see the section *“The Greater Los Angeles dataset”* below). Importantly, the analyses presented here are based on the geographic projection of species’ niches, as opposed to the environmental niche space, making them suitable for inferring landscape-level impacts on species’ distributions (Grether et al., 2024). The existing habitat suitability surfaces are used to generate ROC curves and their AUC scores, importantly, by comparing values drawn separately from urban and non-urban locations. This is in contrast to the prevalent application and interpretation of AUC as a metric of model fit, based on comparing habitat suitability values at presence and absence locations (Merow et al. 2013, Araújo et al. 2019, Valavi et al. 2022). By substituting the locations from which habitat suitability values are drawn (presences=urban and absences=non-urban), the resulting AUC values are effectively converted into a metric that summarizes the disparity of suitable habitat distribution, or habitat disparity (HD) in short (see section “*Using the ROC curve and its AUC as a metric of habitat disparity (HD)*” below). To demonstrate the versatility of this approach, I calculate HD_URBAN_ for all 1,023 species and compare values quantitatively to those expected for different population ecological response types, e.g., urban avoider, urban utilizer, and urban dweller species (Blair 1996, Fischer et al. 2015), stemming from a virtual species modeling approach (Hirzel et al. 2001, Meynard et al. 2019). Lastly, I validate the performance of HD_URBAN_ by comparing native and non-native species, by testing for spatiotemporal robustness across species using Blair’s dataset (1996), and by testing the sensitivity to temporal changes within a species known to decline in urban areas, but not in adjacent wildlands.

### The Greater Los Angeles dataset

This provides an overview of the taxonomic breadth, geographical setting, and modeling framework employed to parameterize the habitat suitability models of the published dataset of 1,023 species in Greater Los Angeles (Beninde et al., 2023), which underlies the analyses presented here. All species have at least one fully terrestrial life stage and represent all higher taxonomic groups: plants (45.5%), arthropods (27.45%), vertebrates (22.2%), fungi (3.2%), molluscs (1.3%), and other taxonomic groups (< 0.3%). The study extent covers 7,797 km² in Greater Los Angeles, fully encompassing the City of Los Angeles, large parts of the Los Angeles-Long Beach-Anaheim, CA Metropolitan Statistical Area, plus parts of adjacent Ventura County in the west, small parts of Riverside and San Bernardino Counties to the east, and into Orange County in the south (Figure 1b). Developed land use types predominate (60.3%), followed by mostly vegetated areas (37.2%), while highly managed, working landscapes, such as agricultural areas, are uncommon (0.7%; based on the National Land Cover Dataset; U.S. Geological Survey 2014). Using the binary U.S. Census Bureau land delineation (U.S. Census Bureau, Population Division 2018; Figure 1c), 65.1% of the study extent is urban and home to a dense human population of 13.4 million (2,640 people/km², calculated based on data from Rose et al. 2017). The remaining 34.9% of the area is non-urban and collectively referred to as wildlands. They entail vast expanses of native vegetation, with little development, including parts or all of the Santa Monica Mountains, Simi Hills, Santa Susana Mountains, Verdugo Hills, Griffith Park, San Gabriel Mountains, Chino Hills, Santa Ana Mountains, and the San Joaquin Hills (Figure 1c). They are sparsely populated, with a modest combined population of 86,700 thousand humans and a density almost two orders of magnitude lower (32 people/km²; calculated based on data from Rose et al., 2017), which frames the extensive urban areas within the study extent. This landscape is uniquely suited to study the response of species to a sharp urban-wildland interface separating an urban megacity from the immediately adjacent wildlands (Figure 1a).

The modeling framework utilized observations of species that were available from the community science platform iNaturalist. Such datasets are commonly referred to as unstructured datasets, which emphasizes their associated sampling biases due to the absence of an underlying (scientifically informed) sampling strategy (Kramer-Schadt et al. 2013, Fourcade et al. 2014). The modeling framework applied by Beninde et al. (2023) employed various approaches to address these causes for potential biases, for example, by thinning occurrence records, scaling background point selection to sampling effort, modeling separately in different extents reflecting land-use types (urban or wildland), null-modeling validation, and using different modeling algorithms to train models. For the analyses presented here, I utilize the set of geographic habitat suitability predictions stemming from the most robust modeling approach. This approach (*i*) uses down-sampled random forest modeling following Valavi et al. (2021) on (*ii*) occurrence records thinned to one record per raster cell, (*iii*) a random background point selection, (*iv*) is performed within the entire study extent (as opposed to separately in urban and wildland habitats), (*v*) for species with a minimum of 25 occurrence records (after the spatial thinning of *ii*), and (*vi*) includes only those models that exceed null model expectations. All 1,023 species fulfilled these requirements.

### Using the ROC curve and its AUC as a metric of habitat disparity (HD)

The Area Under the ROC Curve is a statistical method used to evaluate the performance of a continuous index as a binary classifier, originating in signal detection theory to assess radar detection performance (Peterson et al., 1954). It has since been adopted in various fields for evaluating diagnostic tests and predictive models, including medical sciences, psychology, environmental sciences, and machine learning (Pollack and Decker 1958, Hanley and McNeil 1982, Mason and Graham 2002). It provides a visual and quantitative measure of a model’s ability to discriminate between two classes, with the Area Under the Curve (AUC) representing the model’s overall accuracy. The primary advantages of AUC are that it is scale-free, ranging between 0 and 1, and that it is threshold-independent, providing an easily interpretable and readily comparable summary metric of performance evaluated at different thresholds of the index (Merow et al. 2013, Araújo et al. 2019, Valavi et al. 2022). In species distribution modeling, the AUC is widely used to evaluate a model’s predictive ability in distinguishing between presence and absence (or background) locations of species, with the predicted habitat suitability values serving as the index (Pearce and Ferrier 2000, Araújo et al. 2019, Valavi et al. 2022). Performance is then evaluated by plotting the sensitivity (or true positive rate) and 1-specificity (false positive rate) of correctly classifying presences and absences across different thresholds (e.g., from 2x2 confusion matrices for each threshold), which allows calculating the AUC.

In presence-only modeling (the approach underlying the models used here), absences are unknown (or uncertain) and are replaced with background locations (Merow et al. 2013, Valavi et al. 2022). These can be sampled randomly within the modeling extent or scaled to the distribution of sampling effort in the landscape (Fourcade et al. 2014). In its conventional application to evaluate model performance, an AUC score of 1 indicates a theoretically perfect model, where higher habitat suitability values are observed at all presence locations (the targeted outcome class) with no overlap with lower habitat suitability values at all absence locations (Araújo et al. 2019, Valavi et al. 2022). An intermediate AUC score of 0.5 suggests that the model performs no better than random predictions of presences and absences (note, however, that AUC values of random, or null models, vary around 0.5, mainly as a function of the sample size used to train models; Raes and Steege 2007; Beninde et al. 2023). Consequently, AUC scores reported in species distribution modeling are typically well above 0.5, indicating that models predict presences considerably better than random. Conversely, a theoretical AUC score of 0 indicates that the model performs in the opposite direction of its intended purpose, predicting lower habitat suitability values at all presence locations with no overlap with higher habitat suitability values at absence locations. Therefore, models with AUC values approaching 0 are generally disregarded in species distribution modeling due to their lack of utility in predicting species presence, despite their exceptional discriminatory ability. The discriminatory potential of such models has been explored previously, showing improvements to model fit by inverting predictions of models with AUC<0.5 (Fawcett 2004, Flach and Wu 2005). However, to my knowledge, this property of AUC is rarely reported or utilized. This highlights an unexplored opportunity to utilize AUC in a system where the full breadth of its information content is desirable and meaningful. The question of species responses to a focal habitat type presents such a case where the entire AUC space can be utilized effectively, as all kinds of responses are expected and meaningful.

Implementing the area under the ROC curve (AUC) in this manner requires a binary land classification of the study extent into focal and non-focal habitats (Figure 1c). Here, I classify the study extent into urban and wildland habitats based on US census data (U.S. Census Bureau, Population Division 2018; see section “*The Greater Los Angeles dataset*” above). By calculating AUC using habitat suitability values drawn from urban and wildland locations, the resulting scores reflect the disparity in the distribution of habitat suitability between classes (HD_URBAN_). By defining urban locations as the targeted outcome class (in substitution for presence locations), the metric summarizes an affinity towards the focal urban habitat type. It should be interpreted as follows: HD_URBAN_ of 1 indicates the highest possible urban affinity, characterized by high habitat disparity and higher habitat suitability values at urban locations, which do not overlap with lower habitat suitability values at all wildland locations. Intermediate HD_URBAN_ values ∼0.5 indicate no habitat disparity but indistinguishable habitat suitability values between urban and wildland locations. Conversely, an HDURBAN value of 0 indicates the lowest possible urban affinity, characterized by high habitat disparity but higher habitat suitability values at wildland locations, which do not overlap with lower habitat suitability values at urban locations (see the overview in Figure 2).

**Figure 2:**
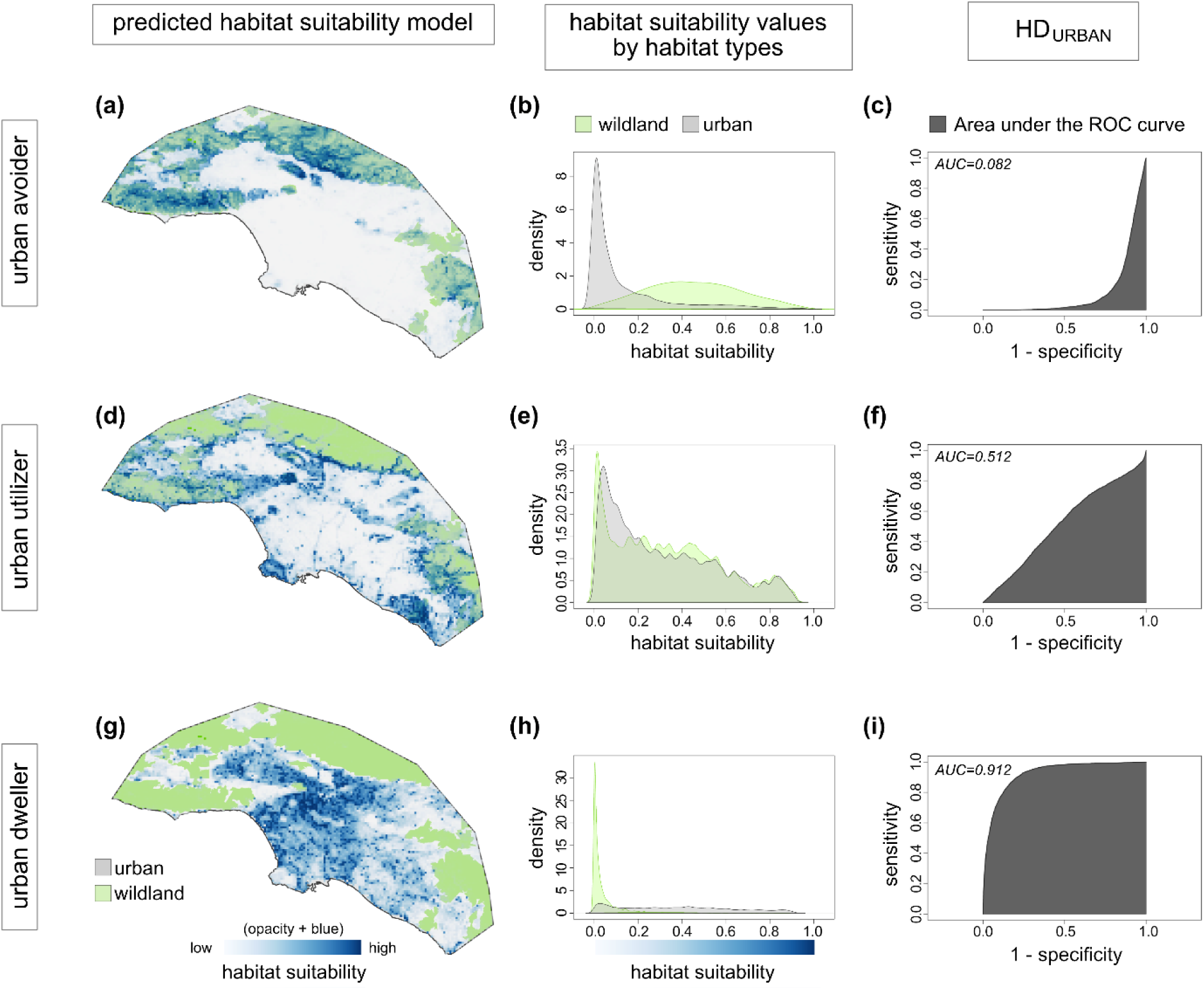
Overview of how HD_URBAN_ is calculated. Examples of virtual urban avoiders (a-c), urban utilizers (d-f), and urban dwellers (g-i) are shown in rows. They illustrate, in columns, the corresponding predicted habitat suitability models, the distributions of habitat suitability values binned by urban and wildland areas, and the resulting HD_URBAN_ scores.

Importantly, HD_URBAN_ measures the degree of disparity in habitat suitability values between classified habitat types, rather than providing a direct measure of habitat preference for a specific intensity of a focal habitat type. While this may seem trivial and conceptually difficult to distinguish, it is a significant difference in measurement. Using the setting of this study as an example, it means that a species with HD_URBAN_ approaching 0 has a much higher habitat suitability in all wildland areas and a much lower suitability in all urban areas; however, this does not necessitate that this species also has the highest suitability in the wildest habitat. And vice versa, a species with HD_URBAN_ approaching 1 may not necessarily have the highest suitability in the most urban habitats.

### Modeling virtual species for urban response types

To facilitate quantitative comparisons between HD_URBAN_ of real species and those expected for the three urban ecological response types, I employed a virtual species modeling approach (Hirzel et al. 2001, Meynard et al. 2019). The virtual species were created reflecting the definitions by Fischer (2015). Fischer defines urban avoider species to “rarely occur in developed areas” and distinguishes urban utilizer and dweller species by varying “relative importance of natural and developed areas to population dynamics”. Further, the persistence of urban dweller species is independent of natural areas, while urban utilizer species still require natural areas for maintaining population dynamics, implying ecological-trap dynamics imposed on these species by urban areas (“breeders […] are present only because of dispersal from adjacent natural areas”; Fischer et al. 2015). Here, I largely adopt Fischer’s definitions, with urban avoider and urban dweller species characterized by opposite responses to urbanization; the former dependent on resources in more natural, wildland areas, and the latter dependent on resources from urban areas. For urban utilizer species, however, I relax the ecological trap dynamic denoted by Fischer and define them as showing no discernible difference (disparity) in suitable habitat between urban and non-urban habitats. This resembles Blair’s initial definition and implies similar breeding success in both habitats in the long term (while acknowledging that ecological traps may govern the dynamics of some species, or temporarily so). Practically, three different virtual species were created, one characteristic for each of the three urban population ecological response types: (a) *urban avoider* virtual species were trained using presence locations exclusively from wildland areas; (b) *urban utilizer* virtual species using presence locations distributed across the entire modeling extent, encompassing both urban and wildland areas; and (c) *urban dweller* virtual species using presence locations exclusively from urban areas. For each virtual species type, 100 down-sampled random forest models were trained using the same modeling framework employed for the real species, and the same environmental predictor data were used (see above for details). The number of presence locations used for training virtual species models was randomly drawn from the number of occurrence records used for training the 1,023 real species (mean N = 146 ± 6.38 SE) and presences were distributed randomly in the landscape given the spatial constraints of each of the response types, i.e., (a)-(c).

### Calculating HD_URBAN_

HD_URBAN_ is calculated from habitat suitability values of 100 randomly drawn raster cells from both urban and wildland areas. The area under the ROC curve (AUC) is then calculated from 2x2 confusion matrices generated across different threshold values of habitat suitability using the evalmod() function in the “precrec” R-package (v0.14.4; Saito and Rehmsmeier 2017), and setting urban values as the targeted outcome class (in substitution of presence values, e.g., when evaluating model-fit). To generate representative AUC estimates, this procedure was repeated 100 times for each species, real and virtual, and the mean was interpreted as their HD_URBAN_ score.

### Validation

Comparing modes of establishment, I tested whether non-native species have higher HD_URBAN_ than native species. Given that urban areas are a frequent pathway for the introduction and subsequent establishment of non-native species (Cadotte et al., 2017; Gaertner et al., 2017; McLean et al., 2017), a higher urban association can be expected for non-native than for native species. This was tested by categorizing all real species with data available (N=996) from Beninde et al. (2023) into non-native (N=705) and native bins (N=291) and testing for differences in HD_URBAN_ using Mood’s Median Test in the “coin” R-package (v1.4-3; Hothorn et al. 2008).

To test for spatiotemporal robustness in urban responses across species, I compared HD_URBAN_ to the urban association ranks provided by Blair (1996) for a set of bird species that both datasets have in common. Blair conducted fieldwork in 1992 and 1993, more than 25 years before the majority of the species observations synthesized by Beninde et al. (2023) were collected (median date of species observations: November 2019). Blair’s field observations took place near Palo Alto, California, USA, setting it more than 450 km apart from all locations in this study (Figure 1). This temporal and spatial difference between the two datasets enables assessment of the spatiotemporal robustness of urban responses across species. Data from Blair’s study were taken from Figure 3 of his publication (1996), which ranks 40 species based on “greatest daily density in business district and greatest daily density in biological preserve.” The business district represents the most urban study site, and the biological preserve represents the most natural. This ordering was compared to the HD_URBAN_ scores generated here using a Spearman rank correlation for the 38 bird species that were common to both datasets.

**Figure 3:**
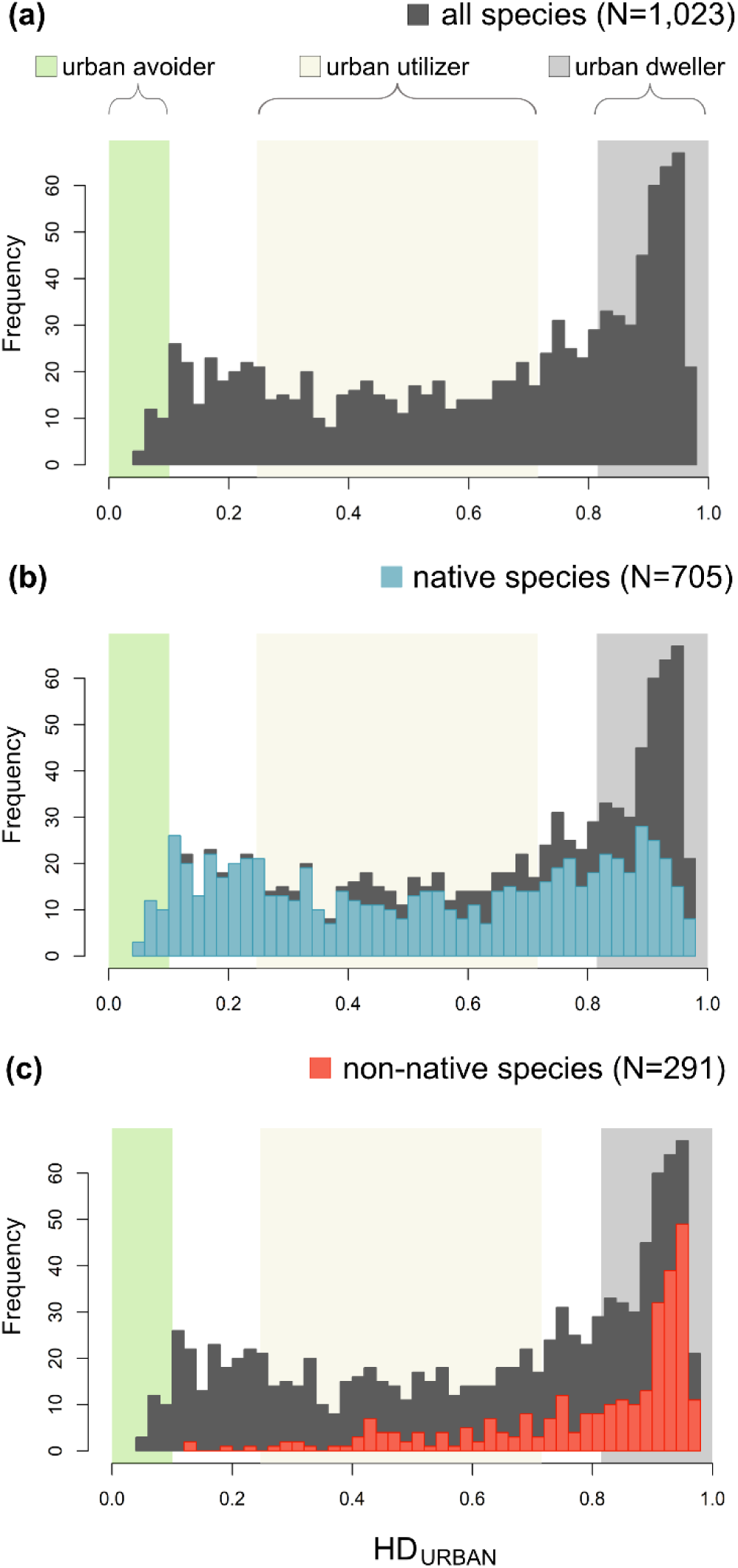
Urban affinity of (a) all 1,023 species, (b) only native, and (c) only non-native species. Expected ranges of HD_URBAN_ for the three urban response types are shown in the background and are based on a virtual species modeling approach.

To assess the sensitivity to detect temporal changes of urban responses within species, I tested for a decrease in HD_URBAN_ calculated for different time intervals for a species known to be declining in urban but not in adjacent wildland areas within the study area. As a case study, I utilize data from the native, Western Black Widow, *Latrodectus hesperus*, and compare it to that of the congeneric, non-native, and recently introduced Brown Widow, *Latrodectus geometrics*. The Western Black Widow has historically been widespread in Southern California, including in urban areas, and it was the only species of *Latrodectus* spider known from Greater Los Angeles until the early 2000s (Vincent et al. 2008). In 2003, the Brown Widow was first reported from within the study extent (Torrance, Los Angeles County; Figure 1c) and subsequently spread rapidly (Vincent et al. 2008). By 2008, it had been recorded in as many as 31 cities in Los Angeles and Orange counties, but remains restricted in distribution to urban areas (Vincent et al. 2008, Vetter et al. 2012, Aragon-Traverso 2021). This rapid spread coincided with a local decline of Western Black Widows in urban areas, a pattern not observed in adjacent wildland areas. Here, Western Black Widows persist while Brown Widows are mostly absent (Vetter et al. 2012, Aragon-Traverso 2021). Depicting these dynamics, HD_URBAN_ for the Western Black Widow is expected to decrease over time within the study area. In contrast, those for the Brown Widow are expected to be stable (and high) since its introduction in 2003. To test this, occurrence records of both species were chronologically ordered, grouped into temporal bins of 25 occurrence records each, and used to train habitat suitability models. This threshold was chosen as a trade-off between the number and quality of models that could be generated from the available observations. Twenty-five occurrence records are a suitable and conservative threshold for the number of occurrences necessary for accurate model training (Proosdij et al., 2016), which is applied to the original dataset used here (Beninde et al., 2023). HD_URBAN_ was then calculated following the same methodology as described above. As observations of both species in the iNaturalist dataset were patchy up until ∼2015, creating datasets of appropriate temporal resolution required including observations available from GBIF (2023, 2024) collected by the Spider Survey of the Los Angeles County Natural History Museum, which started in 2002 (Kempf et al. 2021). Together with the more recent occurrences downloaded directly form the iNaturalist website, this enabled the creation of 25 temporal bins from 625 Western Black Widow observations spanning 2002-2025, and 71 temporal bins from 1,775 Brown Widow observations spanning 2003-2025. Before creating bins, all observations were filtered for spatial accuracy and thinned to one per raster cell and year, following the modeling framework described above.

HD_URBAN_ from each temporal bin served as the response variable in betaregression models with time of observation as the primary predictor variable using the “betareg” R-package (v3.1-4; Zeileis et al. 2021). Time of observations was calculated as the month and year of the median observation per bin (06/2002-04/2025 for Western Black Widow and 11/2009-06/2025 for Brown Widow). As alternative hypotheses, I tested for the effects of seasonality, time span, and geographic space covered by bins. Using the month of the median observation per bin, I assigned one of four seasons: summer (June-August), autumn (September-November), winter (December-February), spring (March-May). Furthermore, as species observations accumulated at different rates over time, temporal bins varied considerably in the number of days between the first and the last occurrence records (ranging from 10-3,905 days, median=53 days). This is particularly well-illustrated by the first temporal bin of both species: The first observation of the Western Black Widow occurred in April 2002, and all 25 observations had accumulated by July 2002. In contrast, observations of the Brown Widow started in August 2003 and required until April 2013 to reach the first 25 observations. Therefore, the time span covered by all observations per bin was included as a predictor in the models. Finally, to address the potential effects of varying geographic coverage per bin, I calculated the area of the minimum convex polygon surrounding all observations per bin using the “terra” R-package (Hijmans, 2021), and included it as a predictor variable in the models. Separate models were fitted for each species, with HD_URBAN_ as the response variable and fixed effects including time of observation, seasonality, time span, and area. Parameter estimates were assessed for significance to determine their importance, and pseudo-R-square values were used to evaluate model fit.

## RESULTS

### HD_URBAN_ of virtual species

The distribution of HD_URBAN_ for virtual urban avoider, urban utilizer, and urban dweller species was non-overlapping (Figure 3). HD_URBAN_ of urban avoider species ranged from 0.055-0.101 (median=0.074), urban utilizer species from 0.247-0.716 (median=0.498), and urban dweller species from 0.815-0.926 (median=0.9).

### HD_URBAN_ of real species

HD_URBAN_ scores of all 1,023 real species ranged from 0.056-0.971 and, overall, leaned more toward higher urban association (median = 0.691; Figure 3a; see https://figshare.com/s/fecc0ef8073cefcfdd1^4^ for HD_URBAN_ of all species).

### Validation

Comparing different modes of establishment, HD_URBAN_ of both native and non-native species were significantly different from normal expectations (Shapiro-Wilk test, *p*<0.001), and native species had significantly lower HD_URBAN_ than non-native species (Mood’s Median Test, *Z* = -11.3, *p*<0.001). Native species exhibited an even distribution of HD_URBAN_ scores (Figure 3b), whereas non-native species scores overlapped more frequently with virtual urban dweller species and not at all with virtual urban avoider species (Figure 3c). Table 1 provides an overview of the overlap in HD_URBAN_ between real species and virtual species, summarized by taxonomic class and categorized by native status.

**Table 1:**
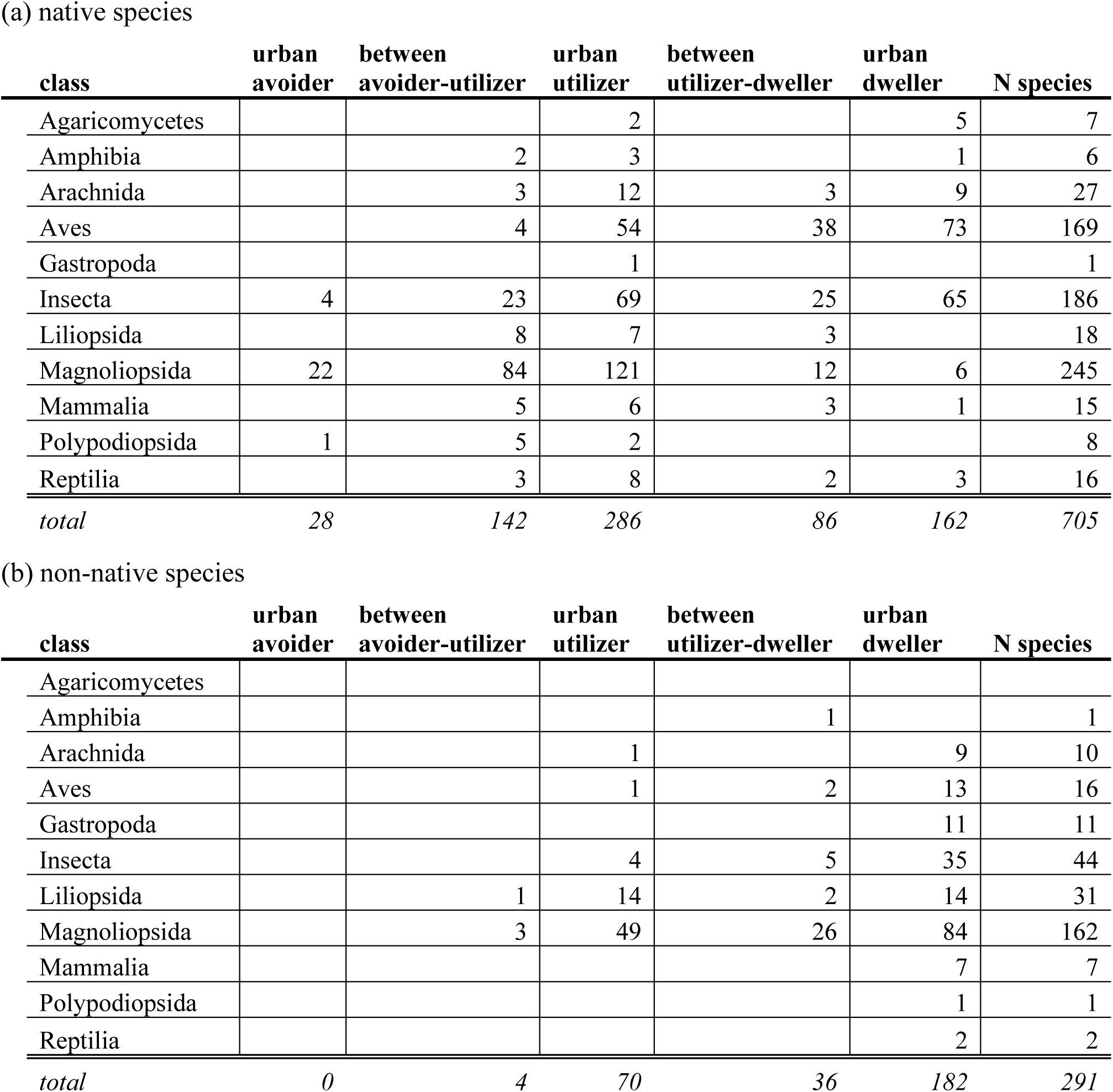
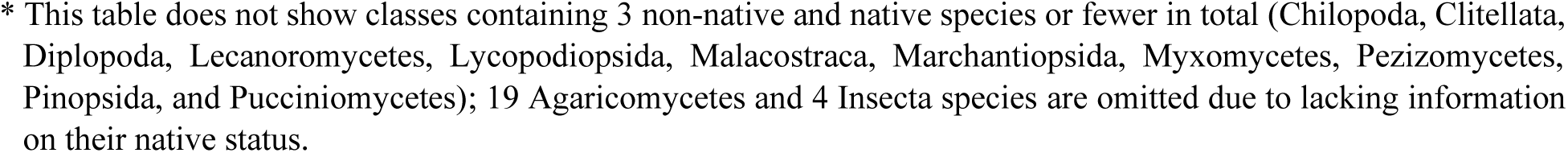
Summary of all real species from the Greater Los Angeles dataset, grouped by their overlap with expected HD_URBAN_ scores per taxonomic class, shown separately for (a) native and (b) non-native species. The expected HD_URBAN_ scores stem from the virtual species modeling approach for each of the urban response types.

Testing for robustness in responses across spatial and temporal scales, the correlation between HD_URBAN_ and Blair’s ranking of 38 bird species was highly significant (Spearman’s rank correlation, S=4202, rho=0.54, p<0.001; Figure 4a).

**Figure 4:**
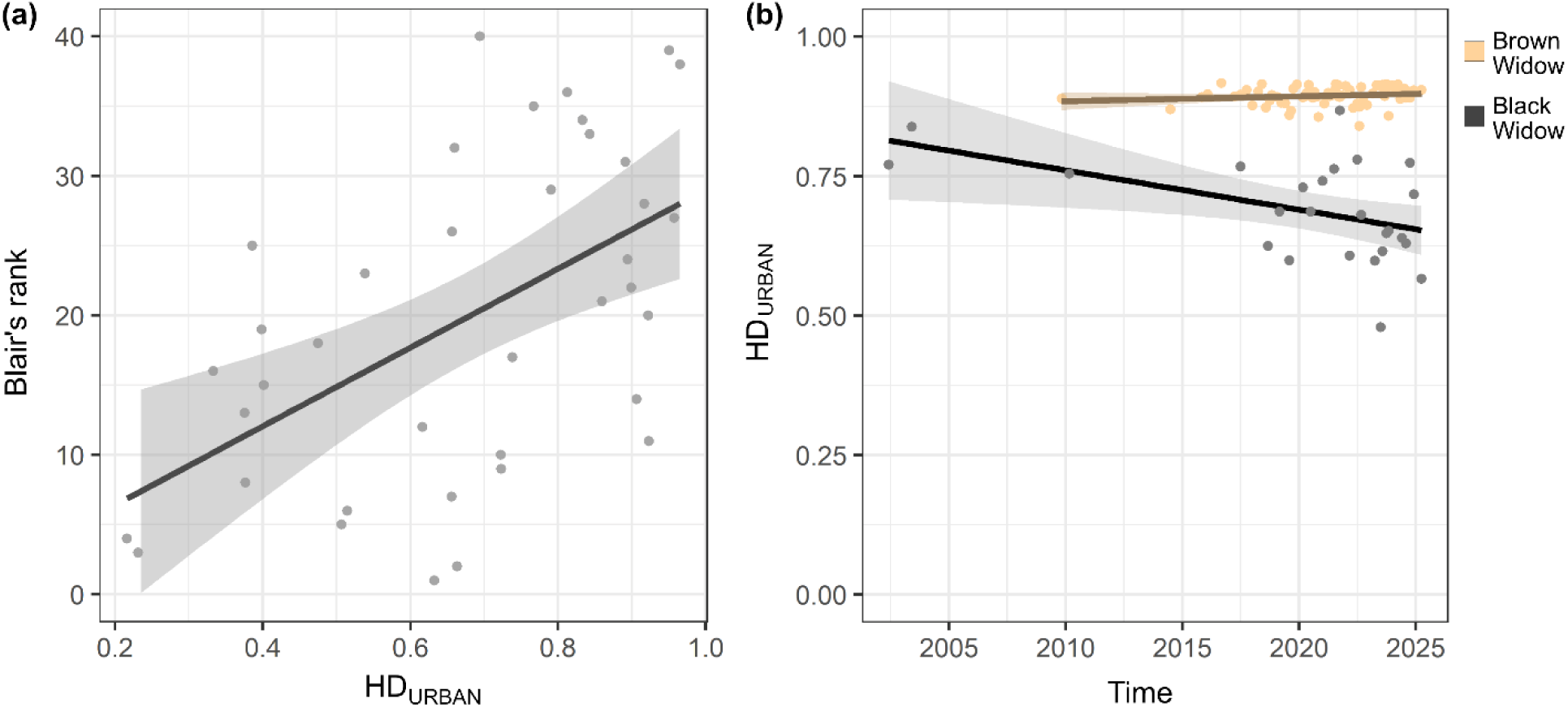
The versatility of HD_URBAN_ is demonstrated (a) by its significant correlation with the ranks of urban associations established for 38 bird species (Blair 1996), showing the robustness of species’ urban responses across space and time, and (b) by its sensitivity to detect changes in urban affinity over time within species, as evidenced by the significant decrease for Western Black Widows, which is being displaced by the Brown Widow from urban Greater Los Angeles since its introduction in 2003.

Assessing sensitivity to detect temporal changes in urban affinity of the Western Black Widows, the time of observation showed a significant, negative effect on HD_URBAN_ (p=0.019) while time span, season, and area remained insignificant (pseudo R² = 0.35; Figure 4b). In Brown Widows, no predictor was significant (all p>0.2), and the total variation explained was low (pseudo R² = 0.04; Figure 4b).

## DISCUSSION

HD_URBAN_ measures the disparity of suitable habitat as a function of the focal urban habitat type, and constitutes an easily interpretable and readily comparable metric of responses to urbanized landscapes, referred to here as urban affinity. This approach relies on geographic predictions of the realized niche of species, obtained from species distribution models (Figure 2), and a novel application of the widely used receiver operating characteristic (ROC) and the area under its curve (AUC). Applied to a dataset of 1,023 species across various plant and animal phyla from the Greater Los Angeles metropolitan area (Figure 1), the results demonstrate that HD_URBAN_ is versatile in application and sensitive to detecting landscape-level, temporal changes in urban affinity of species.

### Validation of HD_URBAN_

As expected, results corroborated significantly lower urban affinity in native than non-native species, with lower HD_URBAN_ in native species (Figure 3). Similar patterns were observed in the Boston metropolitan area, where night-at-light-based urban associations were quantified for 1,004 species across plant and animal taxa (Callaghan et al. 2020b). Given the well-documented and strong association between urbanized landscapes and the introduction pathways of non-native species (Cadotte et al. 2017, Gaertner et al. 2017, McLean et al. 2017), this finding is unsurprising but served as a first test of HD_URBAN_.

Significant robustness of urban responses across species in space and time was demonstrated by comparing HD_URBAN_ to the urban association ranks of 38 bird species established by Blair (1996), assessed 25 years prior and over 450km apart from this study. These findings highlight that, across species, responses to urbanized landscapes can be consistent across urbanized landscapes and temporal scales. For spatial scales, this finding aligns with observations for birds across the Eastern US, where continental levels of Artificial Light At Night (ALAN)-derived urban associations significantly varied with regional urban associations of the same species (Callaghan et al., 2020a). For temporal scales, however, empirical evidence remains more limited. Notably, documented shifts in urban association include two hawk species of the genus *Accipiter*, which have been observed in cities in Europe (Rutz 2008) and Southern California, USA (Cooper et al. 2021), and have both shown increasing urban association over time.

Testing for the sensitivity to detect temporal changes in urban association within species revealed a significant decrease in HD_URBAN_ over time for the Western Black Widow but not for the Brown Widow (Figure 4b). This decline of Black Widows is likely attributed to negative interactions with the congeneric Brown Widow, first detected within the study extent in 2003 and subsequently spreading across urban Greater Los Angeles (Figure 1). Experimental trials have demonstrated preferential predation of Brown Widows on the Southern Black Widow, *Latrodectus mactans* (Coticchio et al. 2023). Assuming similar predation preferences exist for the closely related Western Black Widow studied here, this offers a possible explanation for the observed decline of Western Black Widows in urban areas of Greater Los Angeles and stable population numbers in adjacent wildland areas where Brown Widows are absent (Vincent et al. 2008; Vetter et al. 2012; Aragon-Traverso 2021). The ability of HD_URBAN_ to detect these shifts through time demonstrates its potential for monitoring changes in the urban affinity of species more generally. For example, leveraging the increasing availability of species observation data (Sullivan et al. 2009, Di Cecco et al. 2021), HD_URBAN_ could be deployed to track changes in urban affinity of species globally, e.g., to monitor species targeted by conservation efforts, or to track the succession dynamics and spill-over effects of non-native species (Shaffer 2018; Spear et al., 2018).

### Comparing species to urban population ecological response types using HD_URBAN_

Expanding on these findings, the virtual species modeling approach employed here enables the direct comparison of HD_URBAN_ of real species to that expected of the three urban population ecological response types within the same study extent (Blair 1996, Fischer et al. 2015). Training virtual species for each response type in Greater Los Angeles revealed that all real species overlapping with the HD_URBAN_ of virtual urban avoider species were native. At the same time, many native species also overlap with the high HD_URBAN_ of virtual urban dwellers (Table 1). Amid these are, among birds, the Black Phoebe (*Sayornis nigricans*), Brewer’s blackbird (*Euphagus cyanocephalus*), and Northern mockingbird (*Mimus polyglottos*), and, among insect species, the tobacco hawk moth (*Manduca sexta*), fiery skipper (*Hylephila phyleus*), and figeater beetle (*Cotinis mutabilis*). Native plant species overlapping virtual urban dweller species were less common and included species frequently cultivated as ornamental plants, such as the showy island snapdragon (*Gambelia speciosa*) and apricot mallow (*Sphaeralcea ambigua*; see https://figshare.com/s/fecc0ef8073cefcfdd14 for the HD_URBAN_ of all species).

While the majority of real species overlapping with the HD_URBAN_ of virtual urban-dwelling species are non-native (Figure 3b & 3c), a long tail of low HD_URBAN_ scores for non-native species suggests that some pose a considerable risk for ecological spill-over effects in urban-adjacent wildland areas (Spear et al. 2018). For example, the Spanish broom, *Spartium junceum*, and the yellow star-thistle, *Centaurea solstitialis*, received among the lowest HD_URBAN_ scores of any non-native species (HD_URBAN_=0.19 and 0.23, respectively). They are both flowering plants native to the Mediterranean region in southern Europe, but are known for their invasive populations that occur in similar climatic conditions worldwide, including in California (Bossard et al. 2000).

### Future application and refinement of HD_URBAN_

The data presented here demonstrate the need for a continuous metric for measuring urban responses. HD_URBAN_ values for over 25% of the species did not overlap the expected values of any of the virtual species. Instead, their HD_URBAN_ scores fell between those of virtual utilizers and avoiders (N=146) and those of virtual utilizers and dwellers (N=122). This highlights the continuous nature of responses to urbanized landscapes, expanding beyond the dominating three-tiered typology, and the added value of utilizing a meaningful metric that captures these ecological responses.

HD_URBAN_ facilitates direct comparisons across species and urban landscapes, allowing for the quantification of city-specific effects on urban affinity, such as the scale of cities (Uchida et al. 2021, Blumstein et al. 2023), or environmental differences between urban and urban-adjacent areas (MacGregor-Fors et al. 2021). By comparing urban affinity of the same species across various cities, intra-specific variation of responses and its drivers can be quantified more broadly (Fidino et al. 2021, Haight et al, 2023).

Furthermore, gradient analyses are central in ecological research and extend beyond the prominent urban-rural gradient (McDonnell and Pickett 1990, McDonnell and Hahs 2008, Niemelä 2011, Beninde et al. 2015), encompassing various habitat types, configurations, processes, or impacts (Wilson and Keddy 1986, Curtis and Vincent 2005, Tylianakis et al. 2006, Grundel and Pavlovic 2007, Debinski et al. 2013, Hylkema et al. 2015, Lawrence et al. 2018). Future investigations can help determine the robustness of HD_URBAN_ across different modeling choices and ecological contexts.

Several aspects of HD_URBAN_ calculation, however, deserve further attention and potential refinement. For example, while effective in detecting significant temporal decreases in the Western Black Widow (Figure 4b), the results also revealed high variability in HD_URBAN_ across temporal bins. Future studies could explore the role of temporal bin composition by employing alternative methods to create temporal bins. Moreover, it will be interesting to see if the relatively few urban avoider species observed in this study, compared to urban utilizer or dweller species, can be corroborated across cities or if this might be due to the relative proportions of urban and non-urban areas (here 65.1% and 34.9%, respectively). Additionally, HD_URBAN_ was computed at a specific spatial resolution (∼1km²; Beninde et al. 2023), and optimal resolution may vary based on the availability and accuracy of the species observations. Furthermore, this study trained three virtual species, one for each of the three urban response types, ultimately resting on a binary classification of the landscape into urban and non-urban habitats. Such a binary classification of the landscape may be seen as a limitation of this approach. At the same it necessitates applying objective criteria that distinguish between habitats, which is frequently lacking from urban ecological studies and, ultimately, leads to better comparability across studies, enabling generalizations (MacGregor-Fors 2011).

### Caveats of HD_URBAN_

When utilizing and interpreting HD_URBAN_, two primary aspects require consideration: its specificity in capturing responses induced by the focal habitat type and the computational resources needed.

Firstly, the specificity of HD_URBAN_ to measure urban impacts can vary depending on the landscape’s characteristics and the definition of “urban”. This is well exemplified by the varied topography of Greater Los Angeles, where urban areas are situated at considerably lower altitudes compared to the predominantly hilly wildland areas. Consequently, HD_URBAN_ calculated here reflects urban impacts and altitudinal differences. Moreover, the displacement of urban Western Black Widows is likely driven by negative interactions with the non-native Brown Widow, rather than changing environmental associations. These examples illustrate that HD_URBAN_ is specific to the urbanized landscape under scrutiny, which, importantly, also includes all other local abiotic and biotic factors that covary with it. This must be considered and critically evaluated before interpreting and comparing results across species and landscapes.

Secondly, the calculation of HD_URBAN_ demands substantial observational data and computational resources. A conservative minimum number of 25 observations per species was used here as a threshold for modeling. While this allowed modeling for 1,023 species in Greater Los Angeles, species observations for cities worldwide are less encompassing, and, importantly, detecting temporal trends requires a substantially higher number of occurrence records. Computationally, significant advances have shortened the runtime of high-performance species distribution modeling approaches such as Maxent and random forest to minutes (Valavi et al. 2022). Yet, labor-intensive assembly of spatial predictor datasets from various sources and basic coding proficiency are necessary to apply HD_URBAN_ to new landscapes and to standardize it using virtual species for comparability.

### Conclusion

Quantifying the ecological impacts of urbanized landscapes on populations remains challenging. The metric and methodology presented here demonstrate a path toward data-driven, comparative assessments of these impacts, including the detection of temporal changes in urban responses. This highlights the potential of applying HD_URBAN_ to leverage large-scale community science datasets and develop and compare data across space and time. Finally, by modeling virtual species, real species can be quantitatively compared to the three-tiered urban population ecological response types, aiding the communication of research findings.

## ACKNOWLEDGEMENTS

I am especially grateful to Ryan Harrigan (UCLA, USA), Zac MacDonald (UCR, USA), and Jannik Beninde (NABU, Germany) for their discussions and helpful comments, which significantly improved this manuscript.

